# CHD8 regulates gut epithelial cell function and affects autism-related behaviours through the gut-brain axis

**DOI:** 10.1101/2021.10.02.462735

**Authors:** Ipsita Chatterjee, Dimitry Getselter, Nasreen Ghaneem, Shai Bel, Evan Elliott

## Abstract

Autism spectrum disorder (ASD) is a neurodevelopmental disorder characterized by early onset deficits in social behavior and repetitive behavior. Chromodomain helicase DNA binding protein (CHD8) is one of the genes with the strongest association to autism. Alongside with the core symptoms of ASD, individuals with ASD are reported to have gastrointestinal (GI) problems, and a majority of individuals with CHD8 mutations display GI problems. However, the relationship between autism related genes, such as CHD8, gastrointestinal function, and autism related behaviours are yet very unclear. In the current study, we found that mice haploinsufficient for CHD8 have leaky gut, a dysregulated transcriptome in gut epithelial cells, decreased gut tuft cells and goblet cells, and an increase in microbial load. Specific deletion of CHD8 in gut epithelial cells induced an increase in anxiety-related behaviours in, a phenotype that is often observed in autism and full body knockdown of CHD8, in addition to decreased tuft cells. In addition, antibiotic treatment of CHD8 haploinsufficient mice attenuates sociability deficits. Therefore, the current study determines a pathway for autism-related GI deficits, and how these deficits may play a direct role in the development of autism-related behaviours.

## Introduction

Autism spectrum disorder (ASD) is a neurodevelopmental disorder characterized by early onset deficits in social behaviour and repetitive behaviour(Xu et al., 2018). More than 100 genes have been strongly associated with ASD(Betancur, 2011), while over 800 risk alleles have been identified in sequence-based studies, showing the heterogeneity and complexity of the disorder(Platt et al., 2017). Chromodomain helicase DNA binding protein (CHD8) is one of the genes with the strongest association with autism(O ’ Roak et al., 2012). Several studies have reported various severe mutations in CHD8 gene associated with autism(Bernier et al., 2014; Iossifov et al., 2014; Stessman et al., 2017; Stolerman et al., 2016). Chd8 is a chromatin remodelling factor (Marfella and Imbalzano, 2007) which binds to β catenin(Nishiyama et al., 2012), is located on 14q11.2 and is a regulator of *wnt* signalling pathway(Durak et al., 2016).

Alongside with the core symptoms of ASD, patients have been reported to have other comorbidities. Individuals with ASD are reported to have gastrointestinal (GI) problems(Buie et al., 2010), which includes constipation, bloating, abdominal pain, and diarrhea (Jolanta Wasilewska and Klukowski, 2015). According to varius studies the prevalence of constipation in ASD varies from 20%-33.9% (Oh and Cheon, 2020). Increase in GI problems has also been associated with the severity of ASD (Mayer et al., 2014; Wang et al., 2011). A tight junction barrier protein zonulin, which can affect intestinal permeability, has been found to be increased in children with ASD(Esnafoglu et al., 2017). On the other hand, many studies have shown difference in gut microbiome population in ASD patients(Kang et al., 2018; Rose et al., 2018; Strati et al., 2017) and in animal models of ASD(Sauer et al., 2019).

80% of the patients with CHD8 mutations have also been reported to have gastrointestinal problem, 60% of them reported specific problem such as constipation. Bernier et al, reported that after disruption of CHD8 in the zebrafish have slower GI motility and reduced number of post mitotic enteric neurons(Bernier et al., 2014).

However not much is known about the gut dysfunction in CHD8 mutation and its connection with ASD. It is not clear how a mutation related to ASD may affect gut function. It is further not clear if the gut function may be related to the autism-associated behavioural changes. To gain insight into these questions, here we have used a mice model with heterozygote knockout of CHD8 large isoform to study the gut brain axis. Deletion of both alleles has been reported to be lethal. Katayama et al., had showed that CHD8L^+/-^ mice shows greater brain weight and brain volume than the control mice, consistent with macrocephaly of ASD patients with CHD8 mutation. CHD8L+/- mice showed anxiety like behaviour. They also showed deficits in social interaction, the duration of active contact was reduced in the mutant mice than the controls. Mutant mice did not show any difference in the memory. Additionally, mutant mice had a shorter intestine and tendency of slower motility(Katayama et al., 2016a).

However, the molecular underlining of the GI changes have not been studied. Also, the connection between these and ASD related behaviour remains unknown. In the current study, we have characterized changes in the gut functional and molecular biology of the CHD8L^+/-^ mice and have found evidence that changes in gut biology may play a direct role in the anxiety-like effects of CHD8 haploinsufficiency.

## Results

To gain insight into gut function, intestinal permeability was determined in the CHD8^+/-^ and wild type mice. These CHD8 +/- mice have previously been shown to display autism-related behaviour. First, we performed immunohistochemistry to determine presence of CHD8 in wild type (WT) and CHD8^+/-^ mice gut (Fig. 1A). Higher intestinal permeability has been reported in 36.7% of ASD patients compared to normal subjects (4.8%) (De Magistris et al., 2010). To observe intestinal permeability, mice were orally treated with FITC-dextran, and plasma levels of FITC Dextran was checked at two hours and six hours after gavage. Levels of FITC dextran in the plasma of CHD8^+/-^ mice were increased six hours after gavage, compared to wild type control (Fig. 1B). This suggests increased gut permeability in the CHD8 ^+/-^ mice.

**Figure 1:**
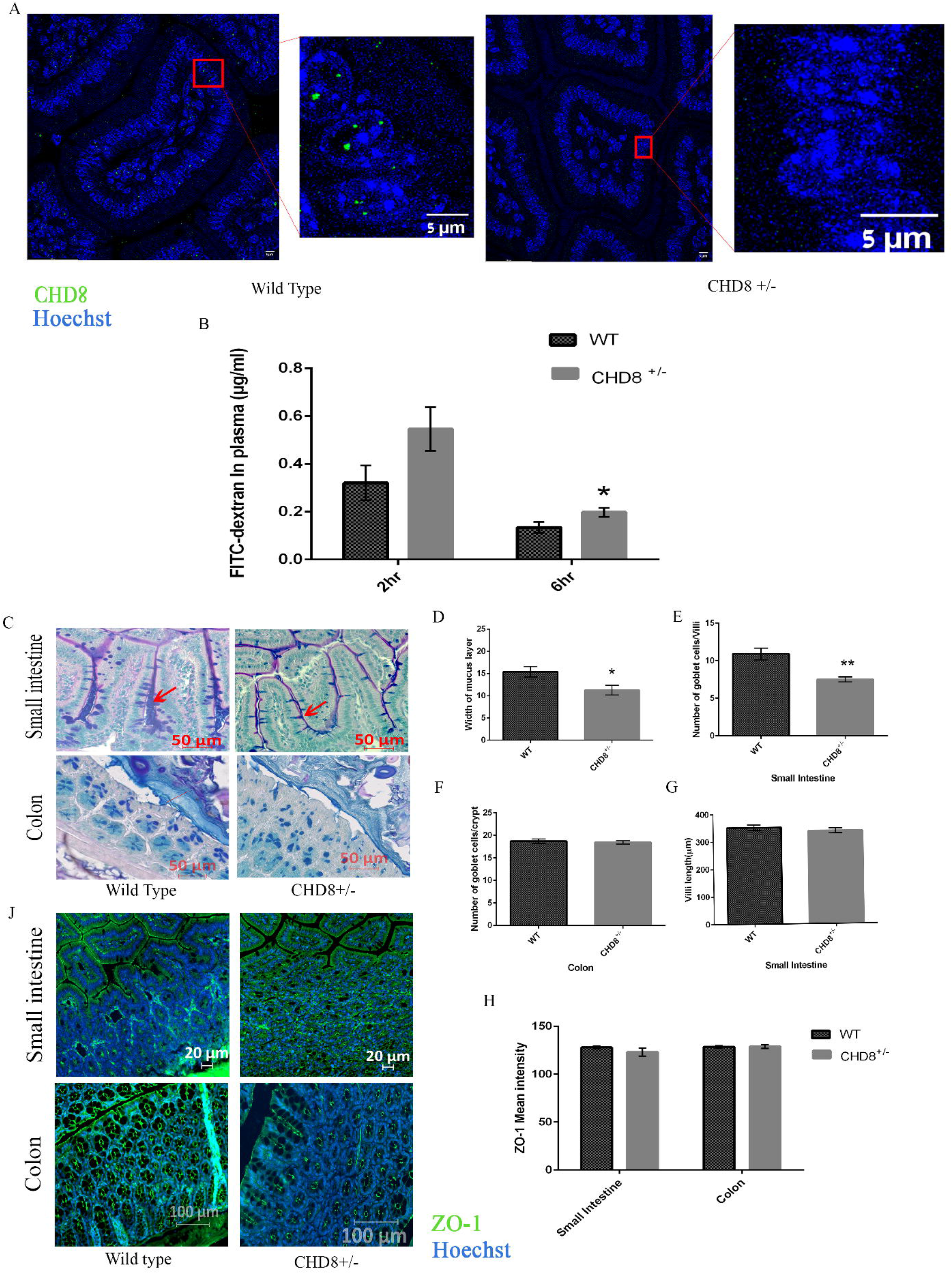
**A**.Representative image of CHD8 staining in Wild type (WT) and CHD8^+/-^ mice. Red box indicates the zoomed area. CHD8 staining is in green and Hoechst staining is in blue. **B**. Intestinal permeability in wild type (WT) and CHD8^+/-^ mice after 2 hr and 6 hr; intestinal permeability was assessed by measuring FITC-dextran level in the plasma after gavage; n=15 in CHD8^+/-^; n=10 in WT. *= p<0.05; Data is presented as mean ± standard error of the mean. **C**. CHD8^+/-^ mice and Wild type mice were stained with periodic acid sciff stain; Representative images are shown, red arrow indicates goblet cells and red line indicated width of mucus layer. **D-G**. Morphological analysis of **D**. Width of mucus layer **E**. goblet cell number/villi **F**. Goblet cell number/crypt, **G**. Villi length reveals a significantly decreased number of goblet cells/villi (Two tailed unpaired T-test; n=8 in WT, n= 6 in CHD8^+/-^, ** = p<0.01) and **F**. significantly decreased mucus layer width (Two tailed unpaired T-test; n=35 in WT, n= 30 in CHD8^+/-^; 5 counting from each animal, *= p<0.05) but not **E**. number of goblet cell/ crypt in colon (p=0.6) and **G**. Villi Length in small intestine (p=>0.05) **J-H**. Immunohistochemistry was performed on 5 mice per group in colon, n=5 in WT, n= 4 in CHD8^+/-^ small intestine, **J**. Representative image of Z0-1 staining in small intestine and colon; **H**. ZO-1 mean intenstity

### Altered gut morphology in CHD8^+/-^ mice

The colonic mucus layer is the main barrier which prevents luminal antigens from coming in contact with host tissues. Thus, we determined whether the elevated gut permeability in CHD8^+/-^ was due to changes in the gut mucus layer. To analyze the gut morphology, we stained the small intestine and colon of mice with AB-PAS staining (Fig 1C), which is mainly used to detect polysaccharides such as glycogen, glycoproteins, glycolipids and mucins in tissue (GOMORI, 1952). Width of mucus layer was decreased in CHD8^+/-^ mice colon (Fig. 1D). In addition, goblet cell number, which produces mucus, was decreased in small intestine (Fig. 1E) although changes were not detected in the colon (Fig. 1F). To understand if these changes could also be detected at earlier developmental time points, we analysed intestines from four weeks old mice. However, there was no differences in mucus layer width or goblet cells between genotypes at the age of four weeks (supplementary figure 1). In addition, we tested for changes in tight junction morphology in eight weeks old mice by staining for the tight junction marker Zo-1. We found no difference in morphology or intensity of Zo-1 staining in CHD8^+/-^ animals, compared to wild types (Fig. 1H). These results suggest that goblet cell and mucus layer decreases may lead to the increased gut permeability in the CHD8^+/-^ mice.

### Transcriptome analysis of gut epithelial cells of CHD8+/- mice

Since CHD8 is a chromatin binding protein, it is likely that CHD8 affects gene transcription in the gut, therefore we performed whole transcriptome sequencing in wild type and CHD8^+/-^ gut epithelial cells. Epithelial cells were extracted from the gut of wild type and haploinsufficient mice, followed by RNA extraction, and whole transcriptome sequencing. Interestingly, over 900 genes were differentially expressed in gut epithelial cells between CHD8^+/-^ and their WT controls. 581 genes were downregulated and 339 genes were upregulated in the CHD^+/-^ gut epithelial cells (Fig. 2A,B). Gene ontology analysis found that downregulated genes were enriched for markers of tuft cells, a subtype of gut epithelial cells, while upregulated genes were enriched for markers of immune response and immune cells (Fig. 2C,D). Biological processes that were enriched in the downregulated genes are involved in mitochondrial function while upregulated genes are enriched for cell cycle related genes (Fig. 2E,F). Considering the strong enrichment of downregulated genes for tuft cell markers, we verified this finding by comparing our downregulated genes to a recently published list of subtypes of tuft cells identified by single cell sequencing. Interestingly, 22 of our downregulated genes are amongst 25 of type 2 tuft cell markers (Haber et al., 2017). We further verified our RNA-seq analysis by real time PCR. We verified a significant decrease in various tuft cell markers (Fig 2G). Tuft cells are known to induce type 2 immune response and induce cells to become goblet cells (Ting and von Moltke, 2019). Therefore, decrease in tuft cells are consistent with the decrease in goblet cells and mucus layer. In addition, RNA-analysis revealed that two of the most upregulated genes were key antimicrobial peptides, Reg3β, Reg3γ. We further verified the increase of these antimicrobial peptides in real time PCR analysis (Fig 2H).

**Figure 2.**
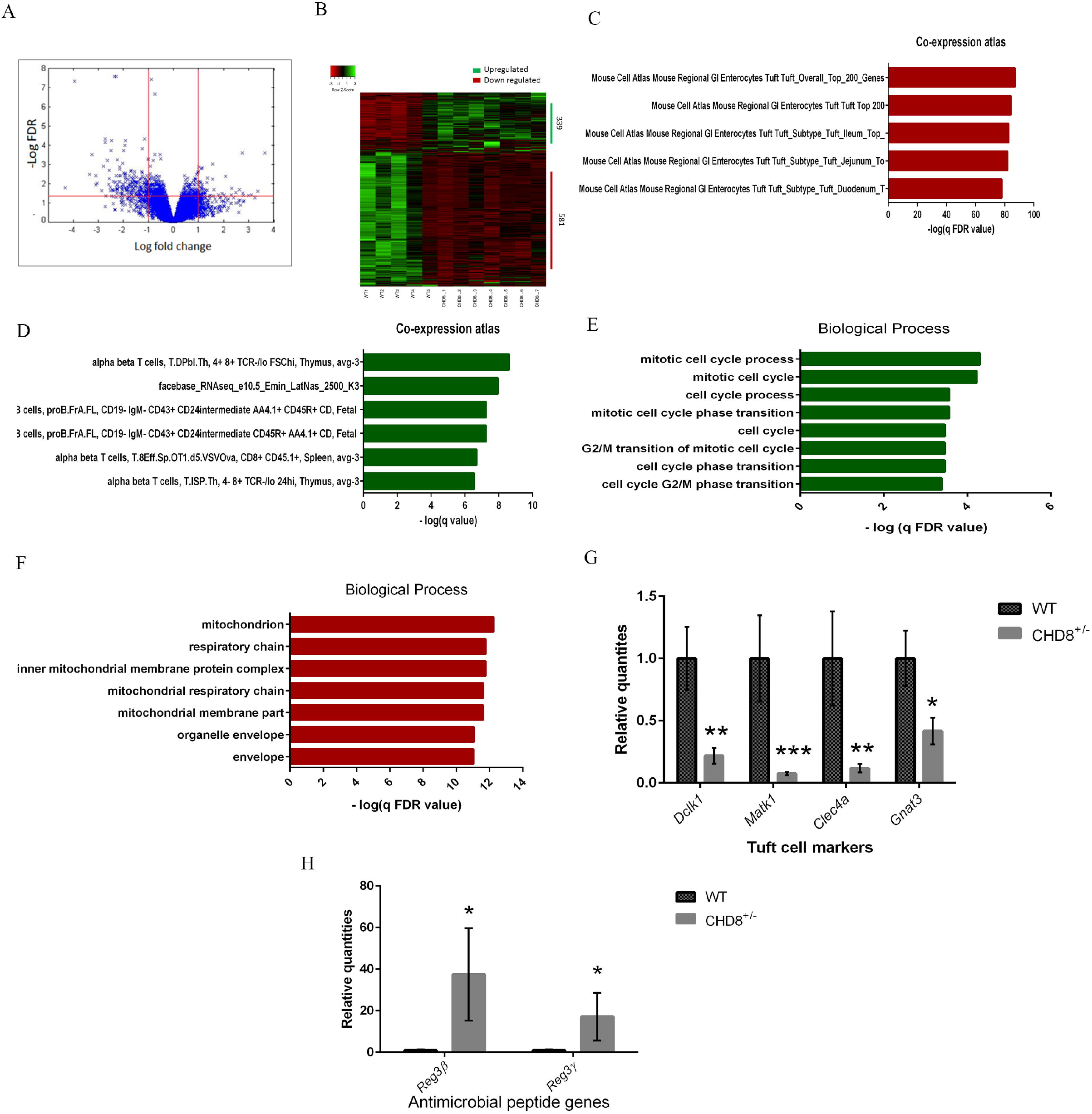
**A**.Volcano plot illustrating the distribution of differential expressed genes in CHD8^+/-^ gut epithelial cells. Each point represents genes in CHD8^+/-^ mice plotted against the level of statistical significance (−log10 adjusted p value) and fold-change (log2 (CHD8^+/-^ vs. WT). **B**. Heatmap showing the 920 differentially expressed genes in CHD8^+/-^ mice gut epithelial cell compared to WT littermates (FDR adjusted p value ≤ 0.05). **C-F**. <u>Gene Ontology</u> analysis of differentially expressed genes. C. Coexpression analysis of downregulated genes D. Co-expression analysis of upregulated genes. E. Biological processes enriched in upregulated genes. F. Biological processes enriched in downregulated genes. G. Real time PCR for tuft cell markers. DCLK1(**=p<0.01), Matk1(***=p<0.001), Clec4a(**=p<0.01), Gnat3(*=p<0.05) (Two tailed t-test unpaired, n=5 in WT, n= 7 in CHD8^+/-^). H. Real time PCR for antimicrobial peptide genes Reg3γ(*=p=0.05), reg3β (*p<0.05) (Two tailed t-test unpaired, n=5 in WT, n= 7 in CHD8^+/-^).

The decreased transcription of tuft cell markers may represent a decrease in tuft cells. To determine if there is a change in the number of tuft cells in the gut of CHD8^+/-^ mice, we performed immunohistochemistry against DCLK1, a marker for tuft cells (Fig 3A). We discovered a decrease in number of DCLK1 positive cells in the small intestine of CHD8^+/-^ mice (fig 3B). No change was observed in the colon (Fig 3B).

**Figure 3.**
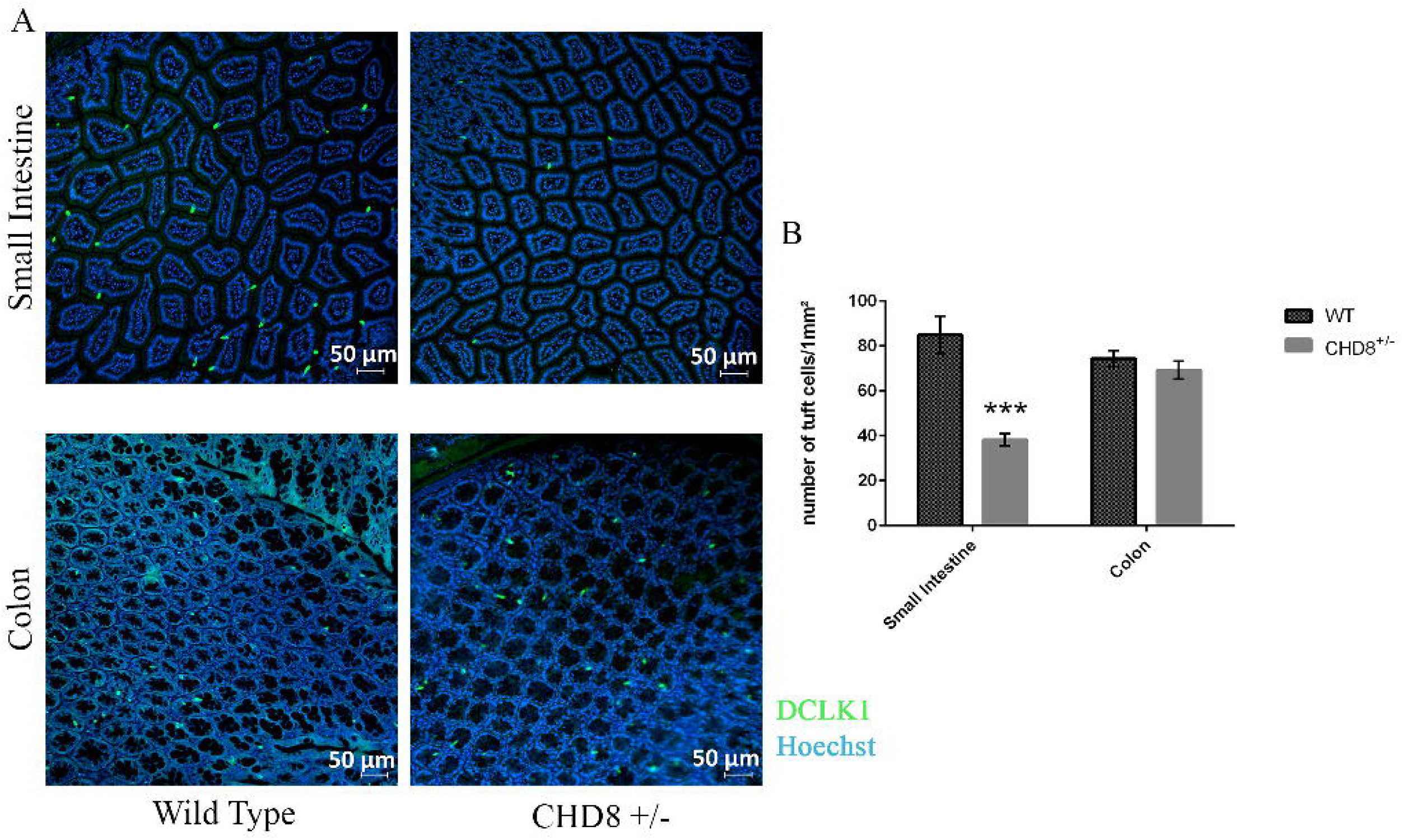
A. Immunohistochemistry was performed on n=4 in WT, n= 5 in CHD8^+/-^ small intestine and colon. Representative pictures are shown here. DCLK1 is stained with green and blue indicates Hoechst staining. B. Number of tuft cells/1 mm^2^ significantly reduced in small intestine of CHD8^+/-^ mice compared to WT (Two tailed unpaired T-test, ***=p<0.001).

### The microbiome in CHD8_+/-_ mice is altered

Considering the possible role of microbiome in autism and the increase in expression of antimicrobial peptides, we determined the microbiome profile in the colon of the CHD8^+/-^ mice and wild types by 16S sequencing analysis. To check the microbiome in this study, Stool was collected directly from the small intestine and colon of 8 weeks old CHD8L^+/-^ mice and their wild type controls.

At first, we checked for total bacterial load in small intestine and colon of these mice by doing real time PCR using primer for 16S as previously described (Nadkarni et al., 2002). In CHD8^+/-^ mice colon, there was a significant increase in overall bacterial load (Fig 4A). 16S sequencing revealed that alpha diversity was increased in the CHD8^+/-^ mice colon (Fig 4B). Beta diversity was not significantly changed between genotype, as visualized in figure 4C. Therefore, CHD8^+/-^ mice display an increase in bacterial load, alpha diversity.

**Figure 4.**
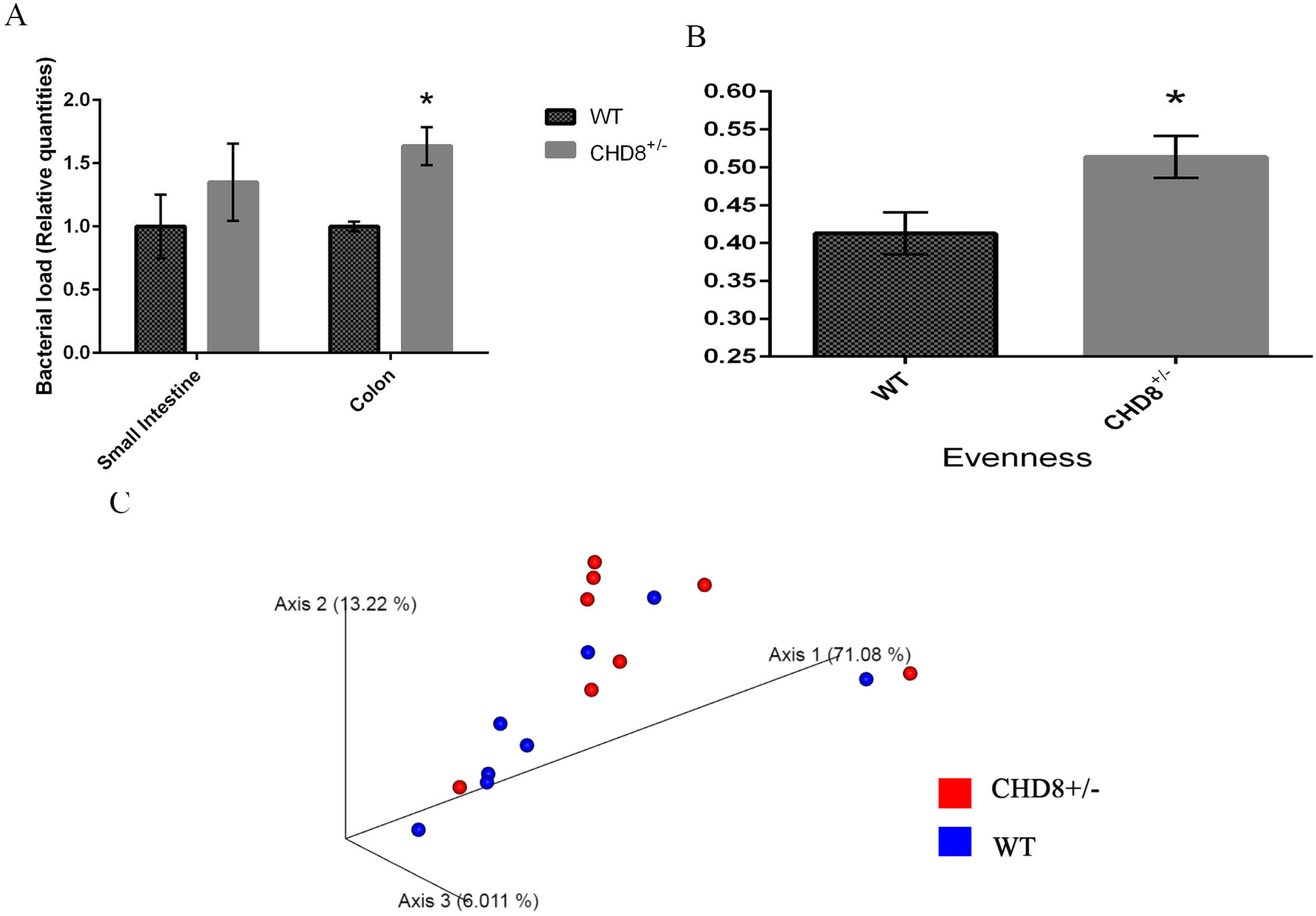
**A**. Relative Bacterial load in small intestine and colon (* = p<0.05) of CHD8^+/-^, WT mice. Two tailed unpaired T-test, n=8 each group. **B**. Measures of alpha diversity; significantly upregulated evenness in CHD8^+/-^ mice compared to WT (Kruskal-wallis pairwise analysis, n=8 per group, *=p<0.05).**C**. Weighted UniFrac-based principal coordinates analysis (PCoA) plot used to visualize microbial communities of all CHD8^+/-^ and WT mice.

### Generation of gut epithelial cell specific CHD8 haploinsufficiency

Considering the complex gut phenotype of CHD8^+/-^ mice, we decided to explore if gut dysfunction can play a role in autism related behavior. To answer this question, CHD8 was specifically knocked out from gut epithelial cells using cre-lox system, with the expression of Cre under the control of gut epithelial-specific promotor Villin. In all experimentation, we compared between haploinsufficient gut epithelial knockdown (Villin-cre/CHD8flx +/-) to wild type littermate controls. Knockdown was validated by immunostaining of the gut in CHD8^ΔIEC^and WT mice (Fig 5A).

**Figure 5.**
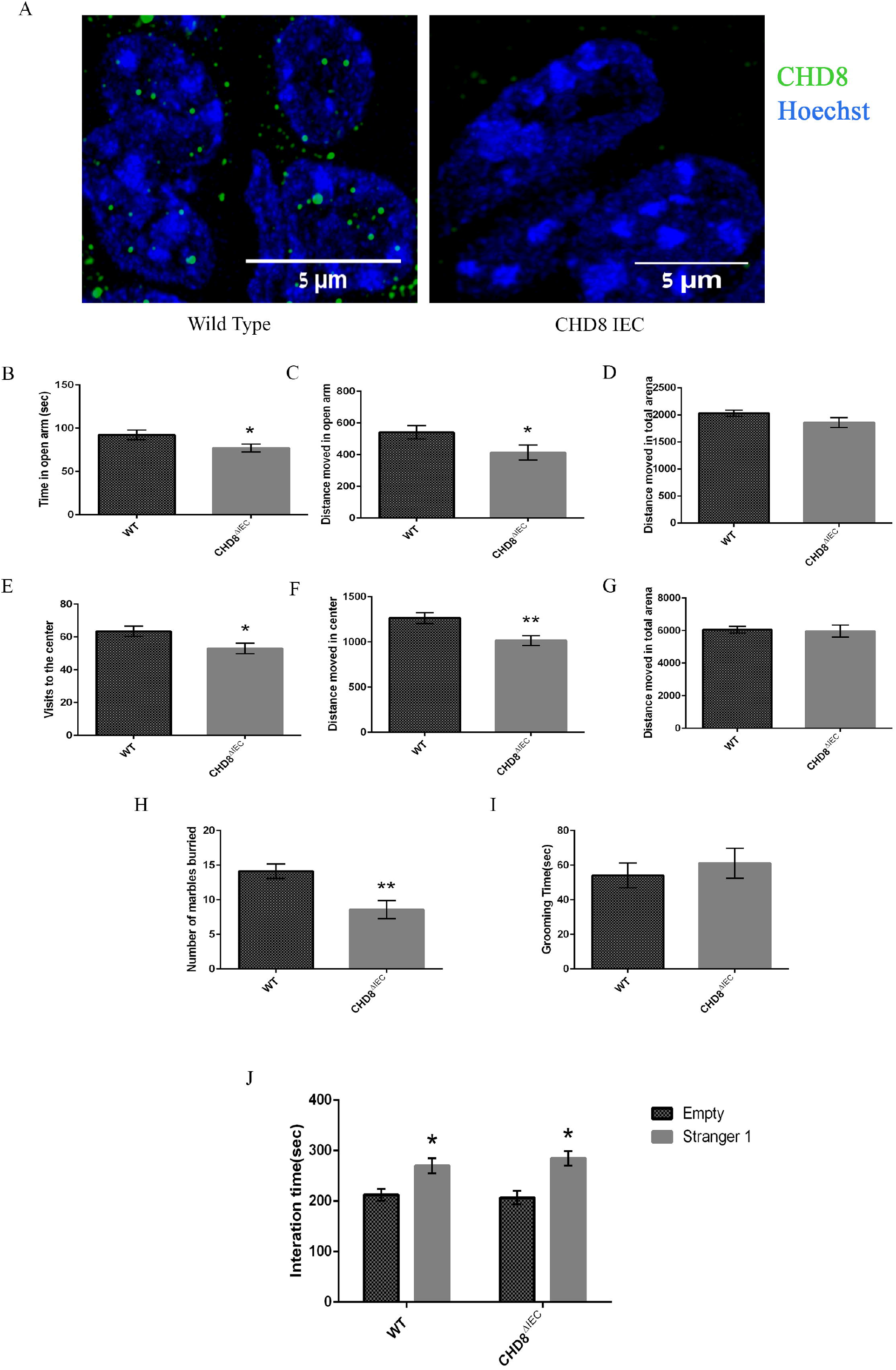
**A**.Representative image of CHD8 staining in Wild type (WT) and CHD8^ΔIEC^ mice. CHD8 staining is in green and blue indicates Hoechst staining. Behavioral test between WT and CHD8 specific KO in epithelial cell mice (CHD8^ΔIEC^). **B-D**. Elevated plus maze; B. Time in open arms; CHD8^ΔIEC^ spend less time. (Two tailed unpaired T test, n=14 in WT, n=11 in CHD8^ΔIEC^, * =p <0.05). **C**. Distance moved in open arms; CHD8^ΔIEC^ moved less distance; (Two tailed unpaired T test, n=14 in WT, n=12 in CHD8^ΔIEC^, * = p<0.05). **D**. Distance moved in arena; No significant difference between groups (p= 0.1). **E-G**. Open field test; **E**. visits to the center; CHD8^ΔIEC^ was significantly reduced than the WT. (Two tailed unpaired T test, n=14 in WT, n=13 in CHD8^ΔIEC^, * =p <0.05). **F**. Distance moved in center; CHD8^ΔIEC^ moved significantly less than the WT. (Two tailed unpaired T test, n=10 in WT, n=13 in CHD8^ΔIEC^, ** = p<0.01). **G**. Distance moved in arena; No significant difference between groups (p= 0.8). **H**. Marble burying test; No of marbles buried by CHD8^ΔIEC^ mice was significantly less than WT. (Two tailed unpaired T test, n=14 in WT, n=12 in CHD8^ΔIEC^, ** = p<0.01). **I**. Grooming time- No significant difference between groups (p= >0.05). **J**. Social preference test-Both WT and CHD8^ΔIEC^ mice showed preference for stranger mice. (Two tailed unpaired T test, n=15 in WT, n=13 in CHD8^ΔIEC^, * represents p=0.005 for WT; p= 0.0005 for CHD8^ΔIEC^).

To check anxiety related behavior in these mice elevated plus maze was done. CHD8^ΔIEC^ mice spent less time in open arms (Fig 5B) and less distance moved in open arms (Fig 5C), suggesting anxiety-like behavior. Consistent with these results, in the open field test, visits to the center (Fig 5E) and distance moved in the center (Fig 5F) was decreased in CHD8^ΔIEC^ mice. Total distance moved was not changed between genotypes (fig 5 D, G). CHD8^ΔIEC^ mice buried less marble than their wild type controls (Fig 5H), which also might suggest anxiety like behavior. No change was observed in the grooming time (Fig 5I), which suggests they do not have changes in repetitive behavior. Three chamber social behavior test was done to explore the social interaction in these mice. They showed no difference compared to their wild controls (Fig 5J). In summary, anxiety-like behavior was specifically induced by knockdown of CHD8 in the gut epithelial cells.

### Reduced Tuft cells in CHD8^ΔIEC^ mice

Considering the decrease of tuft cells quantified in the CHD8 full body haploinsufficient mice, we performed immunohistochemistry on the gut epithelium of the CHD8^ΔIEC^ mice to determine tuft cell numbers. There is a (Fig 6A) significant decrease in DCLK-1 positive tuft cells in both small intestine and colon of 8 weeks old CHD8 ^ΔIEC^ mice (Fig 6A, B). Therefore, gut-specific knockdown of CHD8 induces decreases in tuft cell numbers in the gut epithelium.

**Figure 6.**
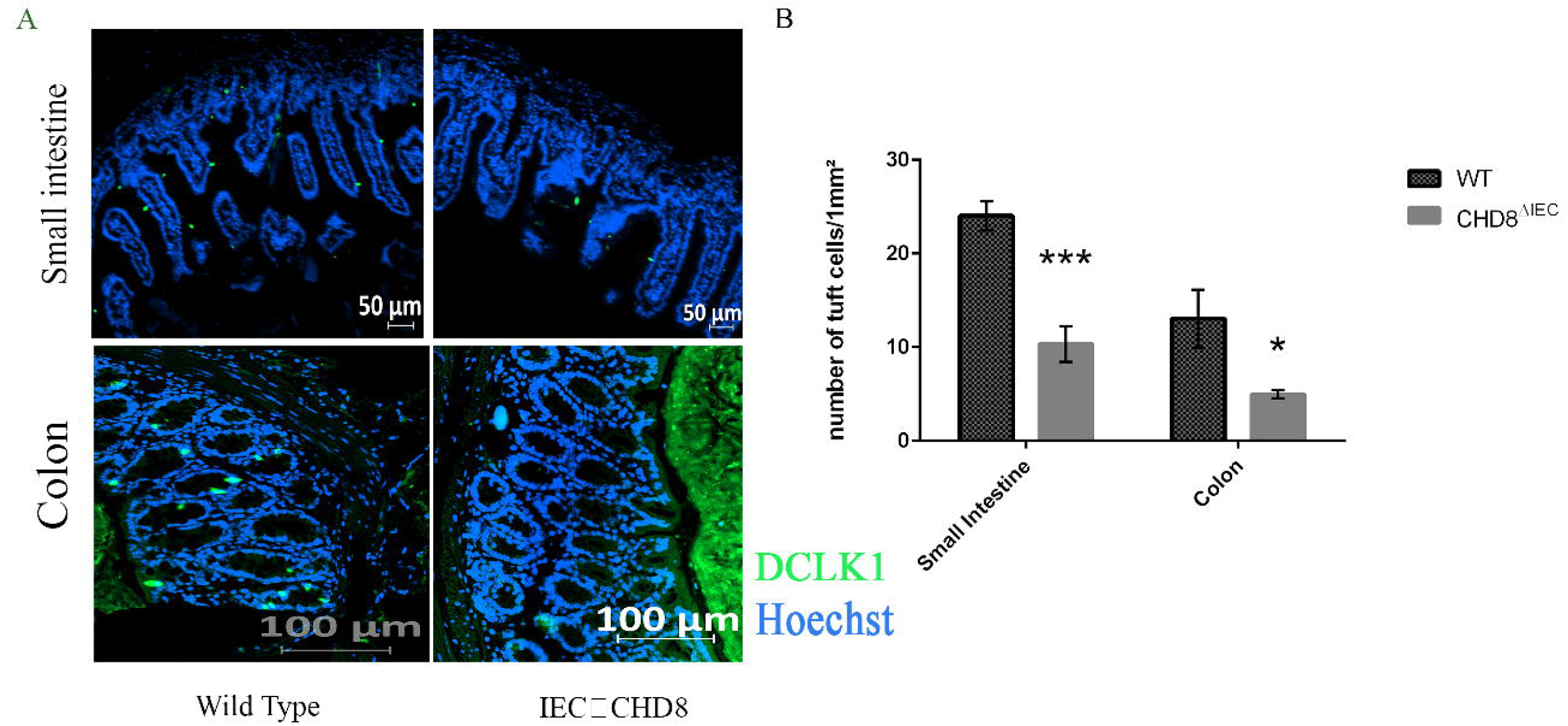
**A-B**. Immunohistochemistry was performed on 5 mice per group. **A**. Representative image of DCLK1 staining in small intestine and colon; **B**. Number of tuft cells/1mm^2^ in small intestine and colon; ***=p<001 *=p<0.05. Two tailed unpaired T-test, n=10 in WT, n=7 in CHD8^ΔIEC^.

### Antibiotic treatment recues social behavior impairments of CHD8^+/-^ mice

Considering that bacterial load was high in CHD8^+/-^ mice, we explored if treating these mice with antibiotics can attenuate their autistic like behavior. 5 week old CHD8^+/-^ mice and their WT controls were given a combination of ciprofloxacin (0.04gl-1), metronidazole (0.2gl-1) and vancomycin (0.1 gl-1) in their drinking water for 3 weeks, followed by behavioral phenotyping. CHD8^+/-^ mice and their WT controls with and without antibiotic treatment were checked for bacterial load by using real time PCR with 16S primers. WT mice with antibiotics and CHD8^+/-^ mice with antibiotics showed significant reduction in bacterial load compared to their without antibiotics controls (Fig 7A). In three chamber social interaction test, WT mice shows preference for the stranger mice compared to the empty cage, both before and after the treatment with antibiotic (Fig 7B). CHD8^+/-^ mice showed no preference for stranger or empty chamber, which suggests deficits in social behavior. Interestingly, after treatment with antibiotics, this deficit was attenuated (Fig 7C). In dark light test, CHD8^+/-^ mice spent less time in open arms than their WT controls (Fig 7D), consistent with katayama et al. 2016(Katayama et al., 2016a). After, antibiotic treatment, no difference was observed between CHD8^+/-^ and their WT controls (Fig 7D). No differences between groups was observed in open field (Fig 7E) and elevated plus maze (Fig 7F). Therefore, antibiotic treatment was able to affect some of the phenotypes seen in the CHD8^+/-^ mice, particularly the social phenotype and one indication of anxiety phenotype.

**Figure 7.**
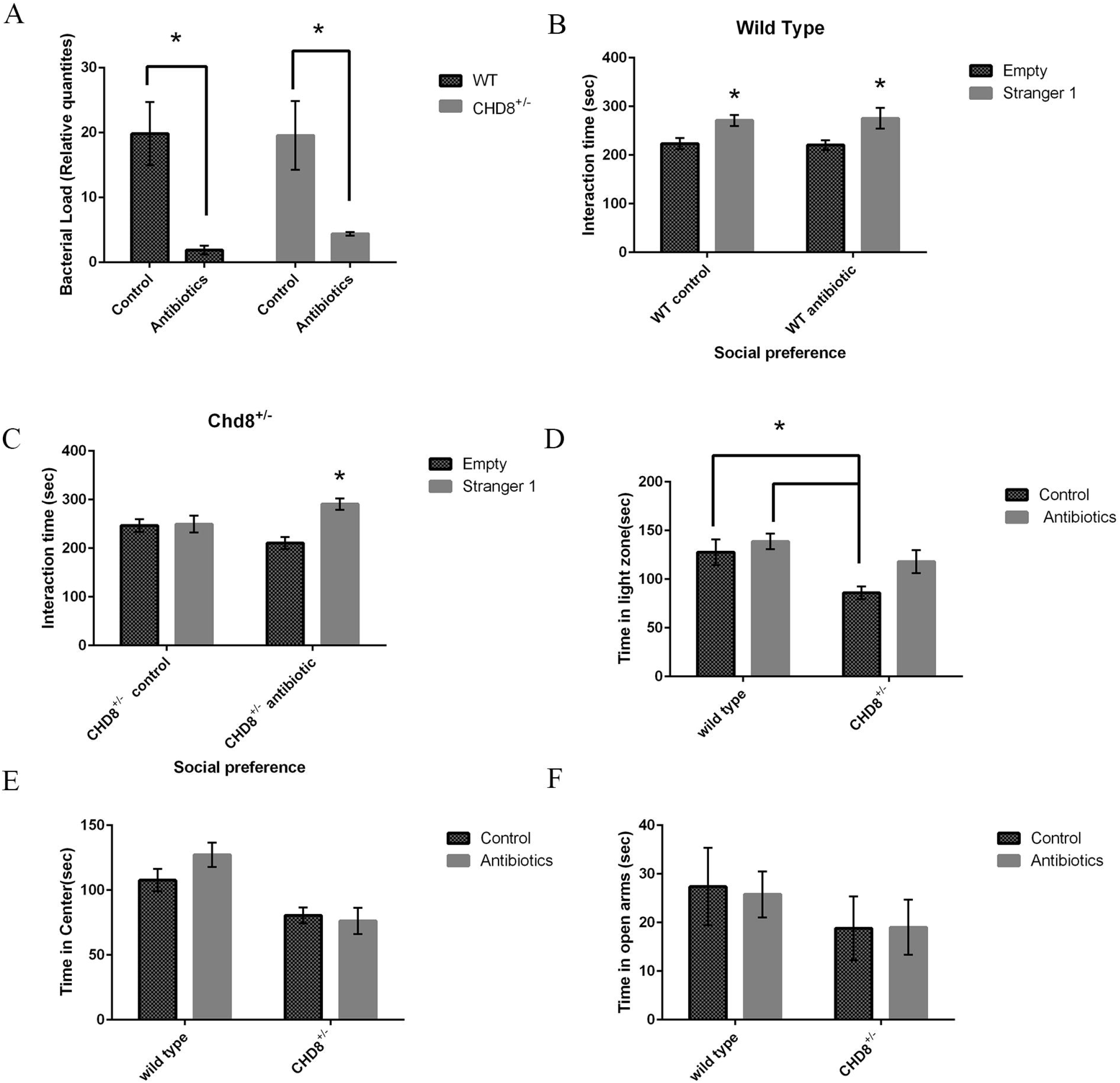
**A**. Real time PCR to check relative abundance of bacterial load after antibiotic treatment; Bacterial load was significantly reduced in WT antibiotics compared to WT control and in CHD8^+/-^ antibiotics compared to CHD8 controls. (Two way anova; n=8 each group, post-hoc Tukey tests * =<0.05). **B-C**. Social preference test. **B**. WT control (n=10) and antibiotics (n=12) mice shows preference towards Stranger mice (Two-way ANOVA, * post-hoc Tukey tests * =<0.05). **C**. CHD8^+/-^ control mice (n=8) shows no preference, CHD8^+/-^ antibiotics (n=9) mice shows preference towards Stranger mice (Two-way ANOVA, post-hoc Tukey tests * =<0.05). **D**. Dark light test-time in light zone; CHD8+/- control mice spend significantly less time in light zone (Two-way ANOVA, post-hoc Tukey tests * =<0.05) than WT control and WT antibiotics mice. CHD8^+/-^ with antibiotics mice had no significant difference than WT control and WT antibiotics mice. (two-way anova, n=11 in WT control; n=12 in WT antibiotics; n=8 in CHD8^+/-^ control; n=9 in CHD8+/- antibiotics.) **E**. Open field test-time in center; **F**. Elevated plus maze-time in open arms; No difference was observed between groups.

## Discussion

GI symptoms have been associated with autism patients, and autism severity has been linked to increased probability of having GI symptoms.(Mayer et al., 2014) Individuals with CHD8 mutations also often display GI symptoms(Bernier et al., 2014).In a previous publication, CHD8^+/-^ mice were reported to have shorter intestine (Katayama et al., 2016b). However, no further analysis on the GI system was done. Zebrafish with chd8 mutations have been reported to have lower motility(Bernier et al., 2014). In the current study, we aimed to understand in depth the relationship between CHD8 haploinsufficiency and gut dysfunction, and its possible roles in the autism behavioural phenotypes. We found CHD8 happlo-insufficient mice to have higher intestinal permeability which is consisted with patients with autism having higher intestinal permeability(De Magistris et al., 2010). Mucus layer is responsible for maintaining the integrity of intestinal permeability(Herath et al., 2020; Johansson et al., 2011). We found in our study that width of mucus layer is lower in CHD8^+/-^, also the number of goblet cells were reduced in small intestine of 8 weeks old chd8^+/-^ mice. Goblet cells produces mucus(Herath et al., 2020), so lower number of goblet cell can cause lower mucus layer, which in turn could have caused higher intestinal permeability. Alterations in GI morphology is consistent with lower villi length in shank3 knockout mice (Sauer et al., 2019). Recently one study showed that foxp1^+/-^ mice shows reduced length of mucus layer in colon of p12.5 mice(Fröhlich et al., 2019). Higher intestinal permeability could also be caused by LPS accumulation, reduction of the tight junction protein ZO-1(Cani et al., 2008) or changes in gut microbiota and their metabolic products(De Magistris et al., 2010). The gut microbiota can affect intestinal permeability by playing an important role in maintaining epithelial integrity(Hsiao et al., 2013).

In our study, we found Chd8^+/-^ mice have higher bacterial load and alpha diversity in the colon. Alteration of gut microbiota has been widely reported in autism patients(Oh and Cheon, 2020). Also, in different mice models of autism changes in microbiome population has been reported(Coretti et al., 2017; Sauer et al., 2019). Transcriptome analysis and real time PCR revealed Anti-microbial peptides like Reg3β and reg3γ were high in gut epithelial cells of CHD8^+/-^ mice. The increase in antimicrobial peptides may be a compensatory mechanism for higher bacterial load.

In our study, by transcriptome analysis of gut epithelial cells in chd8^+/-^ mice compared to WT, we found upregulated genes were enriched in immune system related genes and downregulated genes were enriched in tuft cell markers. Tuft cells are known to induce type 2 immune reaction and are a source of IL25. Il25 drives feed forward tuft-Group 2 innate lymphoid cells (ILC2) signalling circuit. IL5, IL9, IL13 are produced by ILC2 and causes type 2 inflammation. In our study, number of tuft cell positive cells (marked by DCLK1-tuft cell marker) were reduced in small intestine of 8 weeks old CHD8^+/-^ mice. Reduction of tuft cells might cause reduction of goblet cells, and in turn reduction of mucus layer length in these mice.

One of the major behavioral phenotypes of CHD8^+/-^ mice, reported by Katayama et al. was increased anxiety related behaviour. These mice were reported to spend more time in the border in open field test(Katayama et al., 2016b). Very interestingly, in our study, upon the deletion of chd8 specifically from gut epithelial cells, we found these CHD8^ΔIEC^ mice to replicate the anxiety like behaviour of chd8^+/-^ full body haploinsufficient mice. However, no changes were observed in social interaction. Even though we did not see any changes in GI morphology in these mice, we did find number of tuft cells to be significantly reduced in the small intestine and colon of CHD8^ΔIEC^ mice compared to WT controls. This suggests that tuft cells are playing an important role in the anxiety related behaviour. Further studies would be necessary to understand what tuft cell factors may be related to anxiety-related behaviour in these mice.

Gut microbiota has been associated with ASD related behaviours.(Oh and Cheon, 2020). Germ free (GF) mice has been shown to spent more time with an empty chamber than chamber with novel mice, in three chamber sociability test(Oh and Cheon, 2020). In another report, GF mice has shown to be spending more time with novel mice than controls(Arentsen et al., 2015). Administration of single strain such as *Bacteriodes fragilis* has been reported to rescue lack of social behaviour in mice(Hsiao et al., 2013). Additionally, microbial transfer therapy has been proven to improve ASD related symptoms in patients(Kang et al., 2019). Sandler et al, 2000 has shown that patients with regressive onset of autism has shown short term improvement upon oral Vancomycin treatment.(Sandler et al., 2000). Another study determined that children with ASD diagnosis displayed improved SCERTS Assessment Process Observation (SAP-O) scores 6 months after treatment with antibiotics (amoxicillin, Azithromycin)(Kuhn et al., 2012). Consistent with these results, in our study treating the CHD8^+/-^ mice ameliorated the lack of social behaviour. This suggests changes in the microbiome is playing an important role in maintaining social behaviour.

In conclusion, Haploinsufficiency of CHD8 causes changes in GI morphology, tuft cell numbers, gut permeability, and in bacterial load and alpha diversity. Specific knockout of CHD8 in gut epithelial cells (CHD8 ^ΔIEC^) causes anxiety like behaviour but no change in social behaviour. Treating CHD8^+/-^ mice with antibiotics rescues the lack of social behaviour in these mice. This study suggests that GI abnormalities may play an important part in the symptomology and behavioural phenotypes of ASD.

## Figure legends

**Supplementary Figure1.**

**A**. 4 weeks old CHD8+/- mice and Wild type mice were stained with periodic acid sciff stain; Representative images are shown. **B**. Number of goblet cells/villi (p=>0.05). C. Villi length (p=>0.05)

## Materials and methods

### Mice

All mice were bred and maintained in animal facility of faculty of medicine, Bar Ilan University and experimental procedures were approved by Institute Animal Ethical Committee. All mice in a vivarium at 22°C in a 12-hr light/dark cycle, with food and water available ad libitum. C57BL/6 Chd8L+/- mice were crossed with wild type mice and the offspring having CHD8L^+/-^ were used for the experiments with CHD8^+/-^ mice. To generate gut epithelial cell specific CHD8 knock out, the Villin-Cre line was crossed to CHD8loxp/loxp mice (floxed CHD8). Cre-negative (wild type) and Cre-positive (CHD8 floxed heterozygote) littermate offsprings were used in experiments with CHD8^ΔIEC^ mice. All behavioral tests were performed with 8 to 10 weeks old mice.

### Gut-Permeability assay

8 weeks old Mice were fasted for 6 h, and then administrated 14 ml/kg body weight of phosphate-buffered saline (pH 7.4) containing 22 mg/ml fluorescein isothiocyanate conjugated dextran (FITC-dextran, molecular mass 4.4 kDa; Sigma Chemical, St.) by gavage. A blood sample (150 ml) was obtained in a capillary tube 2 h and 6hr after administration of the markers by orbital retro bulbar puncture. The blood samples were centrifuged (3,000 rpm at 4°C) for 15 min. Plasma (50 ml) was mixed with an equal volume of phosphate-buffered saline (PBS; pH 7.4) and added to a 96-well microplate (black). The concentration of FITC-Dextran was determined by spectrophotometry with an excitation wavelength of 485 nm (20 nm band width) and an emission wavelength of 530 nm (25 nm band width) using serially diluted samples of the marker as standard.

### Histology

Small intestine and colon were excised from 8 weeks old mice, immediately submerged in Ethanol-Carnoy’s Fixative at 4°C for 2 hr and then placed into 100% ethanol and subsequently in 50%, 75% and 100% xylene and then embedded in paraffin. It was then cut into 5 µm sections.

#### Alcian-blue Periodic acid sciff’s reagent (Ab/PAS) staining

The tissue sectioned were deparaffinised and stained with alcian blue (sigma; A5268) for 15 minutes. After washing with distilled water, it was treated with periodic acid (sigma Aldrich; p7875) for 5 mints, followed by washing in distilled water for 3 minutes, stained with schiff’s reagent (Sigma Aldrich, 3952016) for 10 minutes. Then it was washed under running tap water for 5 minutes and nuclei were stained with haematoxylin for 1 minute. Sections were then dipped into acid alcohol and then dehydrated and mounted. Number of goblet cells and length of mucus layer was measured using ZEN Desk software.

### Immuno-staining

Sections were deparaffinised followed by antigen retrieval with Sodium Citrate buffer (pH=6) for 20 minutes at 60°C. After cooling down it was permeabilized in 0.1% Triton X-100 for 10 minutes. Blocking was done in 2% BSA (Calbiochem, 126575) and 1% goat serum in 0.1% Tris buiffer saline with triton-X (TbTx) for 3hr at room temperature. The tissue sections were then subjected to immunofluorescence staining with following anitbodies, CHD8(1:100) antibody (abcam: ab84572), ZO-1 (1:200) antibody (Thermo Fisher Scientific; 40-2200), DCLK1 (1:200) antibody (abcam: ab31704) for 1 h at room temperature and followed by overnight incubation in 4°C. The cells were then washed with cold TbTx three times for 10 min each, and incubated with Alexa 488-labeled anti-rabbit secondary antibody (1:200) (Jackson immune research laboratories 111-545-144) at room temperature for 2.30 hr. Then after washing with TbTx 3 times 10 minutes each, they were stained with Hoechst (sigma) (1:1000) while mounting. The sections were examined by fluorescence microscopy.

### Stool collection, DNA extraction and sequencing of 16s rRNA gene

Mice fecal samples were collected directly from the small intestine and colon by scrapping and stored at -80 °C until further analyses. DNA was isolated using the PureLink™ Microbiome DNA Purification Kit according to the manufacturer’s instructions. The V4 region of bacterial 16S rRNA gene was PCR-amplified using the 515F and 806R primers ([CSL STYLE ERROR: reference with no printed form.]). Forward primers included unique 12-base barcodes in order to tag PCR products from different samples. PCR reaction consisted of PrimeSTAR Max Premix 1x (Takara Bio), 0.4 μM of each primer and 30-100 ng DNA template. Reaction conditions were as following: initial denaturing step for 3 min at 95°C, followed by 30 cycles of 10 s at 95°C, 5 s at 55°C and 5 s at 72°C. PCR reactions were performed in duplicates for each sample, pooled and purified with Agencourt AMPure XP kit (Beckman Coulter). Purified PCR products were quantified using a Qubit dsDNA HS assay kit (Life Technologies) and 50 ng of each sample was pooled for further sequencing on the Illumina MiSeq platform.

### Bioinformatic analyses of 16S rRNA gene sequences

Obtained 16s rRNA sequencing data were analyzed by QIIME 1 pipeline (Caporaso et al., 2010). Alpha diversity (within community diversity) was estimated by Gini coefficient, as a measure of community evenness. Beta diversity (between communities diversity) was calculated using weighted UniFrac distances. The diversity parameters were compared between groups using a nonparametric t-test with Monte Carlo permutations (999) to calculate *p* values, and Benjamini and Hochberg FDR method was used afterwards to correct *p* values for multiple comparisons between different pairs of groups.

### Epithelial Cell extraction from gut

Epithelial cells were isolated as previously described by Zeineldin et al(Zeineldin and Neufeld, 2012). Intestinal pieces were washed with PBS, followed by treatment with 0.04% sodium hypochloride for 15 minutes on ice. Then intestinal pieces were put in solution B (details in supplementary table 1 S1) for 15 minutes. Then solution B was discarded and the pieces were put in PBS followed by vortex for 50 seconds. This was repeated 3 times and then the solution was centrifuged at 1000 g, 10 min at 4°C. Epithelial cells are collected in the pellet.

### RNA sequencing and analysis

Collected pellet were resuspended Buffer RLT for RNA purification using the RNeasy Micro kit (Qiagen 74004) following the standard protocol with on-column DNase digestion. Sequencing libraries were prepared using NEBNext Poly(A) mRNA Magnetic Isolation Module & NEBNext® Ultra™ II RNA Library Prep Kit for Illumina® sequencing. The 75 base pair single end sequencing was carried out on the Nextseq 75SR. Differential gene expression analysis was carried out by using DESeq2 pipeline. Enrichment analyses for the Gene ontology (GO) terms (biological process and co-expression atlas) were performed using online ToppGene Suite software. GO terms were considered to be significant when the Benjamini and Hochberg FDR adjusted p value was below 0.05. Raw data and read count data from this analysis are available at GSE182815.

### Real time PCR

RNA was converted to cDNA using Maxima H minus first strand cDNA synthesis kit with dsDNAse (Thermo Scientific). Real time PCR for tuft cell markers and antimicrobial peptide genes was performed using Fast Start Universal SYBR Green Master (Roche) and ViiA™7 Real-Time PCR System (Life Technologies). PCR consisted of 40 cycles, using melting temperature of 95□°C for ten seconds per cycle, and an annealing temperature of 60□°C of thirty seconds per cycle. Relative quantification by ddCt method was used to measure tuft cell abundance in the gut. The primer sequences used in the reactions are indicated in the Supplementary Table S1.

### Bacterial load

Bacterial load was determined as described previously by Nadkarni et al.(Nadkarni et al., 2002) using Taqman and ViiA™7 Real-Time PCR System (Life Technologies). PCR consisted of 40 cycles, using melting temperasture of 95□°C for 20 seconds and an annealing temperature of 60□°C of twenty seconds per cycle. Relative quantification by ddCt method was used to measure bacterial abundance. The primer and probe sequences used in the reactions are indicated in the Supplementary Table S1.

### Antibiotic Treatment

5 weeks old CHD8+/- mice and their WT controls were given a combination of ciprofloxacin (0.04gl-1), metronidazole (0.2gl-1) and vancomycin (0.1 gl-1) in their drinking water for 3 weeks.

### Behavioral testing

Mice were acclimatized to the room for at least 1 h before commencement of each test. Each test was performed on a separate day, usually with one day rest between each test. A camera films the movement, and the Noldus Software “ethovision” tracks the behavior of the animals.

### Open field test

Mouse is placed in the corner of a plastic square box (50×50 cm) where it moves freely for ten minutes under ∼120LUX of light. During this time, a camera films and tracks the behavior of the animals, including distance traveled.

### Light/dark box test

The mouse is placed in a dark plastic chamber (75×75 cm) with an opening to highly lit chamber (∼1200LUX). The mouse is free to move between the two chambers for five minutes. During this time, a camera films and tracks the behavior of the animals, including where they are found inside the box, velocity, distance traveled, etc

### Elevated plus maze

The mouse is placed in the center of a four arms maze. Each arm is 30 cm in length, and two are closed and two are open. The maze is approximately one meter high. The mouse is free to choose which arm it enters for a five minute period. During this time, a camera films and tracks the behavior of the animals, including where they are found in the maze, velocity, distance traveled, etc.

### Social interaction test

The test took place in a Non-Glare Perspex box (60×40 cm) with two partitions that divide the box to three chambers, left, center and right (20×40 cm). The mouse is placed in the middle chamber for habituation (5 min) when the entry for both side chambers is barred. Test mouse was then allowed to explore the whole arena (10 min), where they freely choose between interacting with a novel mouse in one chamber or stay in an empty chamber (social test). During this time, a camera films and tracks the behavior of the animals, including time spent in each chamber.

### Marble burying test

Repetitive marble burying was measured. The apparatus is a Non-Glare Perspex (20×40 cm). Twenty green glass marbles (15 mm in diameter) were arranged in a 4 × 5 grid that covered 2/3 of the apparatus on top of 5 cm clean bedding. Each mouse was placed in the corner that did not contain the marbles and was given 30 min exploration period, after which the number of marbles buried, was counted. “Buried” was defined as 2/3 covered by bedding. Testing was performed under dim light (25 lux).

### Self-Grooming

Mice are scored for spontaneous self-grooming. Each mouse is placed individually into the open field chamber. After 10 minutes habituation period, each mouse was scored using parameters of cumulative time spent grooming during a 20 minutes session.

## Acknowledgements

This work was supported by a grant from SFARI (645766, E.E).

